# Modelling *TERT* regulation across 19 different cancer types based on the MIPRIP 2.0 gene regulatory network approach

**DOI:** 10.1101/513259

**Authors:** Alexandra M. Poos, Theresa Kordaß, Amol Kolte, Volker Ast, Marcus Oswald, Karsten Rippe, Rainer König

**Affiliations:** Integrated Research and Treatment Center, Center for Sepsis Control and Care (CSCC), Jena University Hospital, D-07747 Jena, Am Klinikum 1, Germany; Network Modeling, Leibniz Institute for Natural Product Research and Infection Biology - Hans Knöll Institute (HKI) Jena, Beutenbergstrasse 11a, 07745 Jena, Germany; Division of Chromatin Networks, German Cancer Research Center (DKFZ) and Bioquant, Im Neuenheimer Feld 267, 69120 Heidelberg, Germany; Faculty of Biosciences, Heidelberg University, Germany; Research Group GMP & T Cell Therapy, German Cancer Research Center (DKFZ), Im Neuenheimer Feld 280, 69120 Heidelberg, Germany

**Keywords:** Mixed Integer linear Programming, gene regulatory networks, transcriptional regulation, telomere maintenance, telomerase, cancer

## Abstract

**Background:** Reactivation of the telomerase reverse transcriptase gene *TERT* is a central feature for the unlimited proliferation potential of the majority of cancers but the underlying regulatory processes are only partly understood.

**Results:** We assembled regulator binding information from different sources to construct a generic human and mouse regulatory network. Advancing our “Mixed Integer linear Programming based Regulatory Interaction Predictor” (MIPRIP) approach, we identified the most common and cancer-type specific regulators of *TERT* across 19 different human cancers. The results were validated by using the well-known *TERT* regulation by the ETS1 transcription factor in a subset of melanomas with mutations in the *TERT* promoter.

**Conclusion:** Our improved MIPRIP2 R-package and the associated generic regulatory networks are freely available at https://github.com/network-modeling/MIPRIP. MIPRIP 2.0 identified both common as well as tumor type specific regulators of *TERT*. The software can be easily applied to transcriptome datasets to predict gene regulation for any gene and disease/condition under investigation.

## Background

Telomere repeats are lost at the 3’-end erosion during replication of linear chromosomes. If the telomeres become critically short senescence or apoptosis are induced. This process can thus act as a barrier towards unlimited proliferation and tumorigenesis [1]. Cancer cells circumvent this constraint by acquiring a telomere maintenance mechanism (TMM) [2]. In most instances they reactivate the reverse transcriptase telomerase via different pathways, which can extend the telomere repeats again [3, 4]. Human telomerase consists of the catalytic subunit *TERT* and the template RNA *TERC* (or hTR) [5]. *TERC* is constitutively expressed while the *TERT* gene is silenced in adult somatic cells [6, 7]. Germ and stem cells [7] as well as most tumor cells [2] express *TERT* so that telomerase is assembled. The mechanism of *TERT* activation in cancer cells appears to be highly variable between different cancer entities and numerous transcription factors (TFs) have been reported to be involved in this process [8]. The core region of the human *TERT* promoter is located between 330 bp upstream and 228 bp downstream of the transcription start site. This region comprises several TF binding sites, including binding sites with GC and E-box motifs [8]. Previous studies showed that *TERT* promoter mutations can induce its expression in cancer cells. *TERT* promoter mutations occur most frequently in bladder cancer (59%), cancers of the central nervous system (43%), melanoma skin cancer (29%) and follicular cell-derived thyroid cancer (10%) [9].

Here, we performed an *in silico* analysis of *TERT* regulation by using our previously developed “Mixed Integer linear Programming based Regulatory Interaction Predictor” (MIPRIP) to predict TFs regulating the gene expression of *TERT*. MIPRIP was developed to identify regulatory interactions that best explain the discrepancy of telomerase transcript levels in *Saccharomyces cerevisiae*. In *S. cerevisiae* we uncovered novel regulators of telomerase expression, several of which affect histone levels or modifications [10]. A variety of other approaches have been developed which integrate regulatory information into an unified model of a gene regulatory network (GRN). Many of them infer TF acitvity using linear regression from gene expression profiles, a pre-defined network of TFs and their target genes [11-13], probabilistic models [14] or a reverse engineering approach that identifies regulator to target gene interactions from the pairwise mutual information of their gene expression pofiles [15]. It is noted that the activity of TFs frequently depends only partially on the gene expression of the TF itself but is rather modulated by post-translational modifications and protein stability. Hence, it is informative to infer the activity of a TF from the expression of its potential target genes [11, 16, 17]. In the present study, we have optimized our MIPRIP software and applied it to gene expression profiles of 19 different cancer types from The Cancer Genome Atlas (TCGA) to identify TFs regulating the *TERT* gene.

## Results

### Transcription factor binding information and network construction

We constructed a generic human regulatory network based on seven different repositories, mainly containing experimental validated binding information from ChIP based assays. In total, the generic network comprises 618,537 interactions of 1,160 regulators and 31,915 target genes. For *TERT*, we identified 75 putative regulators (Table S2) that originated mainly from the manual curated database MetaCore™ (60 out of 75). Our list of *TERT* regulators compares well to the *TERT* regulators described in the review by Ramlee *et al*. [8]. In total, 30 from our identified 75 regulators were also described by Ramlee *et al.* (Fisher’s Exact Test P=6.01E-23) and except of CTCF (Encode) all are listed in MetaCore™. Additionally, we assembled a generic gene regulatory network for mouse containing 93,140 interactions of 976 TFs and 15,728 target genes from three different databases. To focus on more reliable edges, entries were selected based on database reliability and co-occurrences.

### Three different modes of a MIPRIP 2.0 analysis

MIPRIP 2.0 can be used to (i) predict the most important regulators of one group of samples (single-mode), (ii) identify significant regulators being different between two groups of samples (e.g. disease vs. control) (dual-mode) and (iii) can be applied to more than two groups (multi-mode). The newly developed multi-mode implementation is embedded in a statistical analysis pipeline and can be applied to more than two datasets or conditions to identify common but also condition-specific regulators (Fig. 1). Here, we applied the multi-mode MIPRIP 2.0 version to study the regulation of *TERT* across 19 different cancer types (described in the next section and Table S3) and employed the dual-mode to compare the regulation of melanoma samples with and without *TERT* promoter mutation.

**Figure 1.**
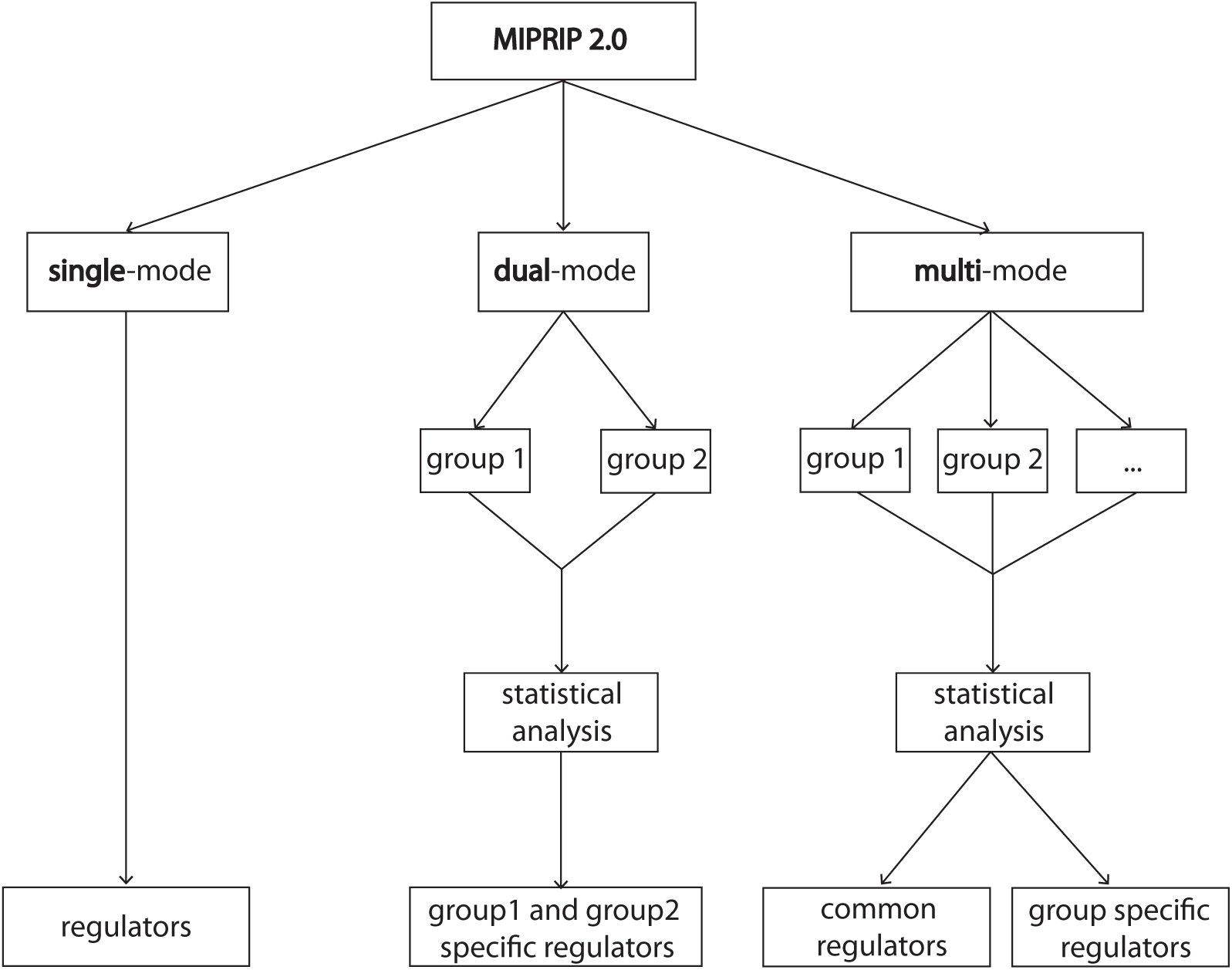
Schematic overview of the workflow. Three different modes are available in MIPRIP 2.0. The single-mode can be used to predict the most relevant regulators of the gene of interest based on a single entity of the disease or condition. The dual-mode compares the regulator predictions of a gene of interest between two different diseases or conditions (e.g. treatment *versus* control). The multi-mode can be used for more than two diseases or conditions to identify the most common and condition specific regulators of the gene of interest.

### Applying MIPRIP 2.0 to identify regulators of *TERT* across different cancers

We selected 19 different cancer types from TCGA for which more than 100 primary tumor samples were available. For each cancer type, we set up a regulatory model for *TERT* by using a ten-times three-fold cross-validation. We calculated different models by restricting the numbers of maximal regulators from 1 up to 10 resulting in 300 models per cancer type. The performance of the models was estimated by the correlation between the predicted and the measured gene expression value (of *TERT* in the expression data). For most of the cancer types, the performance was *r* = 0.4 or better (Fig 2a). For cervical (CESC), ovary (OV) and melanoma skin (SKCM) cancer the performance was distinctively lower. The highest performance was found for testicular germ cell cancer (TGCT) (*r* = 0.75) and thymoma (THYM) (*r* = 0.7), which also showed the highest *TERT* expression over all cancer types (Fig. 2b). The lowest *TERT* expression was found in breast (BRCA), pancreas (PAAD) and prostate (PRAD) cancer. The expression of *TERT* in melanoma skin cancer was comparable to most of the other cancer types, but the performance of the models was the worst (*r* = 0.1) (Fig. 2a). As common regulators of *TERT* across all cancer types, we identified nine regulators: the two paired box proteins PAX5 and PAX8, the E2F factors 2 and 4, AR, BATF, SMARCB1, TAF1 and MXI1 (Table 1). To validate our results *in silico*, we queried Pubmed articles for the identified regulators. We identified 21 out of 1,002 *TERT* articles for our identified regulators which was a significant enrichment for our hits (*p* = 0.013, Tables S4 and S5).

**Table 1.** Predicted *TERT* regulators common to all 19 cancer types

| TF | E-value |
| --- | --- |
| E2F4 | 0 |
| AR | 1.00 E-04 |
| PAX5 | 4.00 E-04 |
| E2F2 | 6.00 E-04 |
| BATF | 3.20 E-03 |
| PAX8 | 6.30 E-03 |
| SMARCB1 | 1.38 E-02 |
| MXI1 | 1.87 E-02 |
| TAF1 | 2.12 E-02 |

**Figure 2.**
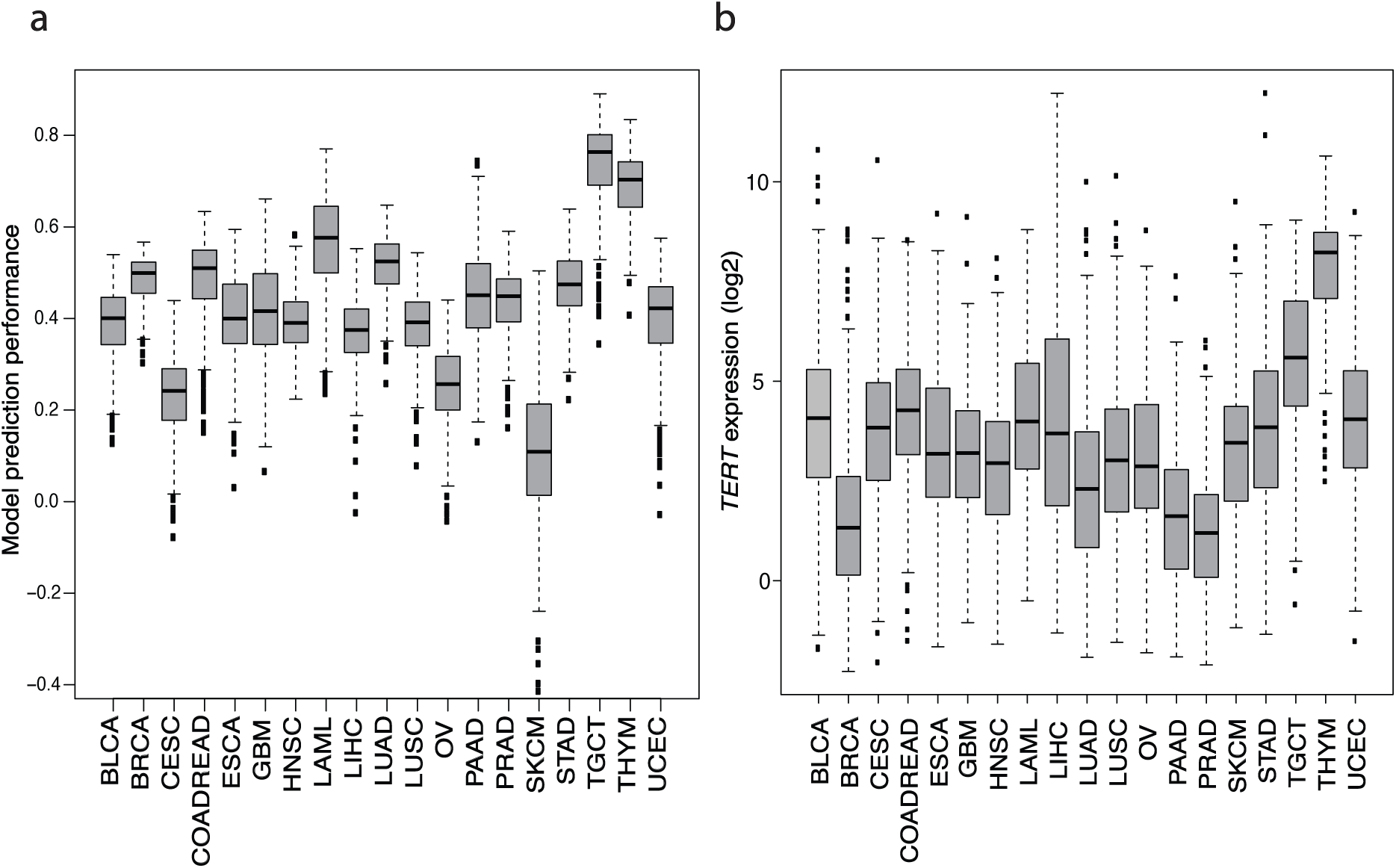
*TERT* expression and prediction performance for the investigated different cancer types. Boxplots for each cancer type of (a) the correlation between predicted and experimental gene expression over all models, and (b) *TERT* expression in each sample.

### Applying the dual-mode MIPRIP analysis to melanoma skin cancer

Melanoma skin cancer was the first cancer type for which a high frequency of *TERT* promoter mutations was discovered, mainly in two hotspot C>T mutations at position 124 bp and 146 bp upstream of the translational start codon [18, 19]. The *TERT* promoter mutation status was available for 115 samples of the melanoma dataset. As described in the previous section, we obtained the lowest performance of our regulatory models for melanoma samples. Considering this and the high rate of *TERT* promoter mutations we divided the dataset into samples with and without *TERT* promoter mutation in order to improve our predictions. We applied the MIPRIP 2.0 dual-mode to the separated datasets. This resulted in a list of 12 and 17 TFs which were significantly more often used in the models for the samples with and without *TERT* promoter mutation. AR, E2F1, JUND and ETS1 were the most significant regulators in the samples with *TERT* promoter mutation, while HMGA2, HIF1, RUNX2 and TAL1 were most significant in the samples without *TERT* promoter mutation (Table 2). To validate that ETS1 is a key regulator in the samples with *TERT* promoter mutation, we investigated published microarray data from experiments in which ETS1 was knocked down in melanoma cells with *TERT* promoter mutation [20]. Indeed, *TERT* expression was lower in the ETS1 knockdown sample compared to controls (fold change: 0.82).

**Table 2.** *TERT* regulators of melanoma samples with (mut) and without (wt) *TERT* promoter mutation

| Regulators in mut | P-value | Regulators in wt | P-value |
| --- | --- | --- | --- |
| AR | 3.97 E-37 | HMGA2 | 1.05 E-16 |
| E2F1 | 3.00 E-29 | HIF.1 | 1.03 E-15 |
| JUND | 2.86 E-25 | RUNX2 | 2.88 E-12 |
| SMARCB1 | 1.85 E-15 | TAL1 | 1.54 E-09 |
| ETS1 | 4.46 E-13 | ESR2 | 3.92 E-09 |
| SIN3AK20 | 1.42 E-06 | AP-2 | 3.28 E-06 |
| REST | 3.64 E-06 | MITF | 2.59 E-05 |
| MAZ | 7.85 E-06 | WT1 | 2.80 E-05 |
| E2F2 | 9.20 E-05 | SMAD3 | 5.76 E-05 |
| TAF1 | 1.36 E-04 | TFAP2D | 2.78 E-04 |
| BCL11A | 4.31 E-04 | PAX8 | 6.04 E-04 |
| MYB | 4.73 E-04 | GRHL2 | 7.16 E-04 |
|  |  | TP53 | 1.21 E-03 |
|  |  | TCF7 | 1.68 E-03 |
|  |  | MZF1 | 3.44 E-03 |
|  |  | TFAP2C | 3.68 E-03 |
|  |  | NR2F2 | 7.96 E-03 |

In summary, splitting up the melanoma dataset into two pre-defined cancer subgroups with and without the *TERT* promoter mutations led to more reliable modelling results (*r* = 0.3). Thus, dual-mode MIPRIP 2.0 is well suited to identify this specific regulation.

### Comparison with ISMARA

We compared our results with results from the well-established tool ISMARA [11]. Similar to MIPRIP, ISMARA identifies the activity of regulators based on their target genes [11]. In contrast to MIPRIP, target genes are inferred from motif binding information, and not directly from ChIP experiments. In ISMARA, TF activities are calculated for each sample alone and then averaged over the samples of each group (*TERT* promoter mutated *versus* wild-type). ISMARA identified twenty TFs for the *TERT* promoter (Supplementary Table S6), but only SIN3A, MAZ and WT1 overlapped with the MIPRIP 2.0 results. Remarkably, ETS1 was not found by ISMARA.

### Availability and Implementation

MIPRIP 2.0 is implemented as a software package in R [21]. It is freely available at our website [22] and on github [23]. MIPRIP 2.0 is platform independent and runs on R version 3.5.1 together with Gurobi version 8.0.1 and the CRAN R package slam.

## Discussion

In the present study we have advanced our software package “Mixed Integer linear Programming based Regulatory Interaction Predictor” (MIPRIP) for application to human and mouse cells. For this, we selected known regulator binding information to construct a generic network linking TFs to their potential target genes. The interactions between TFs and their targets organize as a scale-free network comprising hubs as central regulators [24]. The TFs with the highest score in our generic human regulatory network were MYC (sum of es: 12,607), YY1 (sum of es: 10,642.25) and CTCF (sum of es: 9,574.50). This might reflect the role of these TFs as master regulators that recruit chromatin modifying co-factors and remodel the chromatin structure in the case of MYC [25] or mediate structural interactions between enhancers and promoters as reported for CTCF and YY1 [26].

The MIPRIP 2.0 framework with its new multi-mode was applied to dissect the regulation of the telomerase protein subunit *TERT* across 19 different cancer types, yielding nine TFs being common to *TERT* regulation across all cancer types. These regulators together showed a significant enrichment in Pubmed entries for *TERT.* Five TFs (PAX5, PAX8, AR, E2F2 and E2F4) have been described previously as *TERT* regulators. PAX5 has two and PAX8 four binding sites at the *TERT* transcription start site that induce activation of *TERT* transcription and their function in telomerase regulation has been validated [27, 28]. The androgen receptor (AR) belongs to the class of nuclear receptors and is a repressor of *TERT* expression [29]. The E2F2 and E2F4 factors bind to the E2 recognition motif and are involved in cell cycle processes and DNA damage response [30] and regulate *TERT* transcription in human B-cell lymphoma [31, 32]. In addition to these five TFs, we here identified BATF, SMARCB1, TAF1 and MXI1 as novel *TERT* regulators across cancer entities that to our knowledge, so far, have not been described in the literature as *TERT* regulators. Accordingly, we suggest these as potential candidates for future investigations on the mechanism of *TERT* reactivation in cancer cells.

The best performance of the MIPRIP 2.0 multi-mode analysis was observed for thymoma and testicular germ cell cancer, which showed also the highest *TERT* expression. The worst performance was observed for melanoma skin cancer, even though *TERT* expression was not particularly low. As described in the literature, cutaneous melanoma skin cancer patients have a high rate of *TERT* promoter mutations, being responsible for an upregulation of *TERT* by enabling a further binding site of TFs from the ETS family [18, 19].

Using MIPRIP 2.0 in the dual-mode after dividing the melanoma dataset into cancer samples with and without *TERT* promoter mutation improved the results considerably. We identified ETS1 as a highly significant regulator for *TERT* in tumors with *TERT* promoter mutation. To further validate this finding we analyzed publicly available expression data of an ETS1 siRNA knockdown experiment in a melanoma cell line with *TERT* promoter mutation and found a downregulation of *TERT* compared to controls. In line with this finding, ETS binding together with the activation of the non-canonical NFkB signaling pathway through the co-activator p52 enhances the promoter activity of *TERT* [33].

Besides ETS1, we predicted AR, E2F1 and JUND as the most significant regulators in melanoma patients with a *TERT* promoter mutation. AR and E2F were also predicted as common *TERT* regulators in our multi-mode MIPRIP analysis. A recent study showed that an inhibition of E2F1 leads to increased cell death in melanoma cells, even if they are resistant to BRAF-inhibitors [34]. These results indicate that E2F1 is an interesting therapeutic target for melanoma. According to our predictions, E2F1 regulates samples with a *TERT* promoter mutation. As E2F1 is a *TERT* repressor [30], an inhibition of E2F1 may be more efficient in samples without *TERT* promoter mutation.

For melanoma samples without *TERT* promoter mutation, we predicted HMGA2, HIF1, RUNX2 and TAL1 as the most significant regulators. HMGA2 is a member of the high-mobility group of AT-hook proteins, which are expressed during embryonic development [35] as well as in different tumors (e.g. squamous cell carcinoma and malignant melanoma [36]). While only a few samples showed a *TERT* promoter mutation [36], it is still unclear if there is an association between HMGA2 expression and *TERT* promoter mutations. According to our predictions, we suggest that *TERT* regulation by HMGA2 and *TERT* promoter mutations are mutually exclusive, which has to be validated in future experiments. In this case study, we observed that splitting up the datasets into subtypes led to an increased performance of the regulatory models and was necessary to break down the relevant regulatory processes. Melanoma patients with *TERT* promoter mutation show decreased survival rates [37]. Hence, identifying subtype specific regulatory mechanisms may support risk stratification by employing the identified regulators as biomarkers. In addition, such predictions may pave the way for a personalized therapy by developing drugs specifically interfering with the detected TFs.

Using the specific application of known ETS binding site mutations in the *TERT* promoter of melanoma samples as a case study, we compared the results from MIPRIP 2.0 with ISMARA. The overlap between our results and ISMARA was very low. ISMARA did not retrieve ETS1 as distinct *TERT* regulator of samples with the promoter mutation.

Besides the advancement with the three different modes and the possibility of weighted edges, MIPRIP 2.0 allows to extend the model by including information about gene copy number, DNA methylation, miRNA expression and binding, and additional variables e.g. related to further epigenetic regulation.

## Conclusions

We here introduced our new MIPRIP 2.0 framework and applied it to predict *TERT* regulators in a pan-cancer analysis. Some of the common TFs identified like PAX5, PAX8, AR, E2F2 and E2F4 have been previously described as *TERT* regulators. Others like BATF, SMARCB1, TAF1 and MXI1 are novel. It will be exciting to test experimentally whether they are linked to a TMM phenotype. Furthermore, the predicted *TERT* regulators were compared in melanoma samples with wild-type vs mutated *TERT* promoters. In this manner, we validated that a change of TF targets, in this case for TFs from the ETS family, was captured by MIPRIP 2.0. The software package is available on our website [22] and github [23] together with the generic human or mouse regulatory network and example datasets. It can be applied to a large variety of datasets to investigate the role of TF mediated gene regulation of a gene of interest in the context of diseases or response to a variety of conditions that include treatment.

## Methods

### Gene expression data

We downloaded publicly available transcriptome expression data (RNA-Seq) of all cancer types with more than 100 primary tumor samples from the TCGA Genome Data Analysis Center (GDAC) of the Broad Institute [38]. For these datasets the usage restriction has been lifted according to the TCGA publication guidelines from December 21, 2015 [39]. The pre-processed transcriptomic data with log2 transformed RSEM [40] normalized values were downloaded for 19 different cancer types listed in Table S1. In each cancer type, genes with more than 25% missing entries and low varying genes (standard deviation ≤ 0.5) were filtered out. Furthermore, we performed a z-score transformation for each gene across each cancer dataset.

### Assembling transcription factor binding information into a generic human and mouse gene regulatory network

We assembled a comprehensive set of putative regulators for each gene by compiling TF binding information in human cells from seven different data repositories comprising (i) MetaCore™[41] with annotated “direct”, “indirect” and “unspecific” interactions, (ii) the ChIP Enrichment Analysis (ChEA) database [42], (iii) chromatin immunoprecipitation data from the ENCODE project (http://www.genome.gov/Encode/), (iv) human ChIP-seq and ChIP-ChIP data from hmCHIP [43], (v) experimentally verified interactions from the Human Transcriptional Regulation Interactions database (HTRIdb) [44], (vi) ChIP-seq data for long non-coding RNA and microRNA genes from ChIPBase [45] and (vii) the method of Total Binding Affinity (TBA) [46]. TBA estimates the binding probability of a TF to the whole range of a gene’s promoter. Only TBA values with a stringency cutoff of a score ≥ 1.5 were selected. All these repositories were used to compute the generic network of TFs and their target genes. An interaction between a TF *t* and a target gene *i* was considered if it was listed

i. in MetaCore™and labelled as direct, or listed in Encode,
ii. in at least two out of MetaCore™ (labelled as indirect), ChEA, TBA (score≥1.5) or HTRI, or
iii. in ChIPBase and hmChIP.

The different repositories were not equally incorporated due to the fact, that some repositories were more reliable than others. Because the interactions from MetaCore™based on literature reports and were manually curated, MetaCore’s direct interactions (*MCdir*_*ti*_, activation, inhibition or unspecific) were weighted by a factor of 2, while MetaCore’s indirect interactions (*MCindir*_*ti*_, activation, inhibition or unspecific) were weighted by a factor of 1. Entries from *chea*_*ti*_, *htri*_*ti*_ and *tba*_*ti*_ were also weighted by a factor of 1, interactions from Encode (*enc*_*ti*_) by 0.5. A factor of 0.25 was used for interactions found in hmChIP (*hm*_*ti*_) and ChIPbase (*chip*_*ti*_). This led to the overall edge strength score *es*_*ti*_:

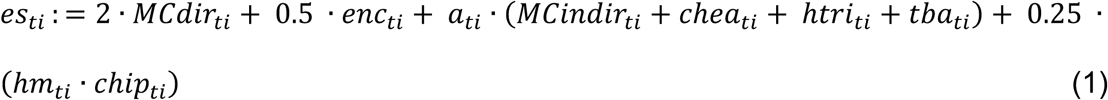

with

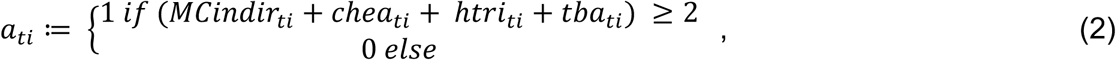

and

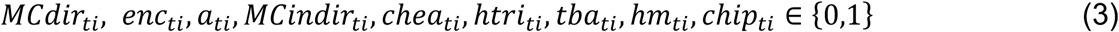

In total the generic network (version 1.0) comprises 618,537 non-zero entries for 1,160 TFs and 31,915 target genes.

Similarly, a comprehensive set of putative regulators for each gene was assembled for mouse by compiling TF binding information from MetaCore™, ChEA and ENCODE containing TF binding information for mouse. Additionally, we added two more databases, ECRBase and TfactS. ECRBase is based on alignments of evolutionary conserved TF binding sites [47]. TfactS [48] contains interaction information inferred from the regulation of TFs from gene expression data of experimentally well-characterized target genes listed in TRED [49], TRRD [50], PAZAR [51] and NFIregulomeDB [52]. Interaction information of TF *t* and target gene *i* from MetaCore™ (MCdir_ti_) labelled as direct was weighted by 2. If an interaction was listed in two out of (a) MetaCore™ indirect (MCindir_ti_), (b) ChEA (chea_ti_) and (c) ECRbase (ecrbase_ti_), it was weighted by 1 (for each source). A listed mouse ENCODE entry (enc_ti_) was weighted by 0.5. The interactions of TfactS (tfacs_ti_) were considered to have weaker evidence and were weighted by 0.25. This led to the overall edge strength score *mes*_*ti*_ for mouse:

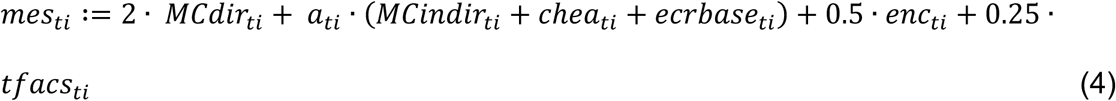

with

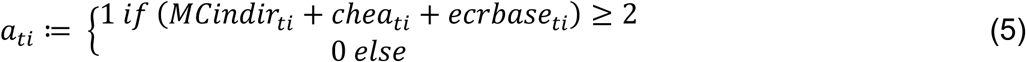

and

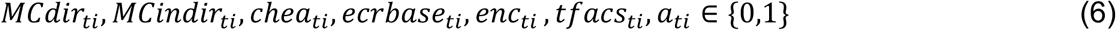

In total the generic mouse network (version 1.0) comprises 93,140 non-zero entries for 976 TFs and 15,728 target genes.

### Modeling *TERT* regulation

We optimized our previously developed “Mixed Integer linear Programming based Regulatory Interaction Predictor” (MIPRIP) software. MIPRIP 2.0 can be used for one set of samples (single-mode), can be applied to compare the regulatory processes between two sets of samples (dual-mode), and for multiple datasets, to identify the most common and condition specific regulators (multi-mode) (Fig. 1). The basic idea of MIPRIP is to identify the most relevant regulators of a particular target gene by predicting the target gene’s expression using a linear model in which the covariates are all potential regulators putatively binding to its promoter. The gene expression value 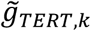 of the *TERT* gene is predicted for each sample by the following model:

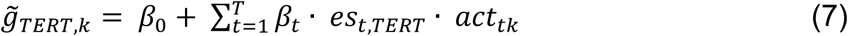

where *β*_*0*_ is an additive offset, *T* the number of all regulators for which *TERT* promoter binding information is available, *β*_*t*_ is the optimization parameter for regulator *t, es*_*ti*_ is the edge strength score between regulator *t* and its putative target gene *i* and *act*_*tk*_ the activity of regulator *t* in sample *k*. If gene *i* was reported to be a target of regulator *t*, the edge weight was higher than 0. Instead of using the gene expression value of a regulator, we calculate an activity value *act*_*tk*_ for each regulator and each sample based on the expression of all its putative target genes *g*_*ik*_ by

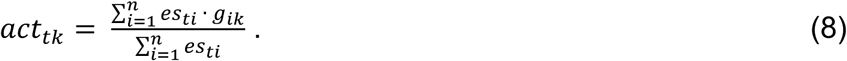

The activity is the cumulative effect of a regulator on all its target genes, normalized by the sum of all target genes. To calculate the activity value, we excluded the expression value of the gene of interest (*TERT*) itself. A linear regression is performed based on Mixed Integer Linear Programming (MILP). MILP has advantages over lasso regression model, as in MILP based regression, the error penalties are linear (L1 regression) and not quadratic which avoids over-emphasizing outliers. Furthermore, MILP enables using binary on-off switches for each beta coefficient to limit the number of beta coefficients [for details, see [17]]. All linear equations are optimized using the Gurobi optimizer [53] (version 6.0-7.01) to minimize the difference between the measured transcript level (from the gene expression matrix) *g*_*i,k*_ and the predicted gene expression 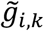 value. This equals to minimizing the error terms *e*_*ik*_ in

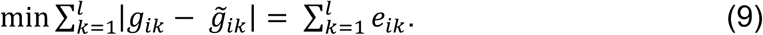

Because MILP cannot handle absolute values, the absolute values were transformed into two inequalities for each gene *i* and sample *k*,

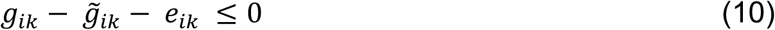

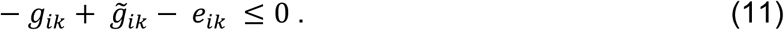

To avoid overfitting, for each dataset we constructed models constraining the number of regulators starting with one regulator up to 10 regulators. A ten-times threefold cross-validation was performed yielding 300 models for each dataset. The correlation between the measured and the predicted gene expression values from the models indicates the prediction performance.

### Single-mode MIPRIP 2.0 analysis

The single-mode analysis was developed to predict a list of regulators best explaining the gene expression profile of the target gene of interest (*TERT*) for all samples of a dataset for a single condition or disease. The single mode has no additional statistical analysis beyond the linear modelling. The results are simply the frequency of regulators over all cross-validation runs, prioritized by their usages.

### Dual-mode MIPRIP 2.0 analysis

As a case study, we applied the dual-mode analysis to the skin cutaneous melanoma (SKCM) dataset. The SKCM dataset was divided into two subgroups based on the *TERT* promoter mutation (based on the analysis of [54]). In total, the status of the *TERT* promoter was available for 115 samples (primary and metastatic samples). One subgroup (n=74) comprised samples with a *TERT* promoter mutation, the other subgroup comprised samples with the according wild-type of the *TERT* promoter (n=41). With these two subgroups we performed a dual-mode analysis by calculating the linear models using the same parameters as described above. Significant regulators between the two subgroups were determined by a two-sided Fisher’s exact test, testing an enrichment of a TF to be in a model of the first or the second condition based on their distribution in the different models, followed by multiple testing correction using the Benjamini-Hochberg method [55]. The stringency cutoff was set to P=0.01.

### Multi-mode MIPRIP 2.0 analysis

The multi-mode analysis was developed to predict (1) a list of regulators best explaining the gene expression of the gene of interest (*TERT*) across all conditions (in our case tumor types), and (2) for each specific condition, in contrast to all other conditions. We prioritized the regulators as follows. For each condition, we listed how often each regulator was selected by the optimizer resulting in a count table. With these distributions we performed a one-sided Wilcoxon Test for each regulator in the list to identify regulators which were selected significantly more often in one of the conditions compared to all other conditions yielding the condition specific. Significance values (p-values) were corrected for multiple testing [55]. To identify the most common *TERT* regulators within all conditions (of all 19 cancer types), a rank product test was performed based on the ranks from the counts of each condition. A permutation-based estimation was used to determine if the rank product value was higher than an observed value from a random distribution. We then counted how often the rank product values in the permutations were below or equal to the observed value, which led to an averaged expected value (E-value) [56].

### Systematic literature query

A Pubmed [57] search was performed for the identified common *TERT* regulators from the multi-mode MIPRIP 2.0 analysis. A combination of all regulator gene symbols from Table 1 was queried together with “TERT” and the terms “telomerase”, “human” and “regulation”: (E2F4 OR AR OR PAX5 OR E2F2 OR BATF OR PAX8 OR SMARCB1 OR MXI1 OR TAF1) AND TERT AND telomerase AND human AND regulation. The received number of articles was compared to a query without the identified regulators, and to the same two queries without the gene symbol of *TERT* yielding the according background counts. A Fisher’s exact test was applied to test if the identified common regulators were significantly more often found together with *TERT* than without *TERT* (Supplementary Table S5).

### TF perturbation experiments

We investigated *TERT* expression upon ETS1 knockdown for validating our result of the dual-mode case study. Wang *et al.* have previously performed siRNA mediated knockdown of 45 TF and signaling molecules in the melanoma cell line A375. The gene expression of cells with the knockdown (1 sample per knockdown), untreated (3 replicates) and siRNA control treated (3 replicates) cells was measured by microarrays (Affymetrix GeneChip Human Genome U133 Plus 2.0) 48 h after transfection [20]. RMA-normalized expression data of the perturbation experiments was downloaded from Gene Expression Omnibus (GSE31534). Affy probe-ids were mapped to gene symbols using BioMart [58] and multiple affy probe-ids for the same gene were averaged. A fold change was calculated for *TERT* upon ETS1 knockdown compared to control.

### Comparison of MIPRIP 2.0 with ISMARA

We compared our MIPRIP 2.0 results with the results from the “Integrated Motif Activity Response Analysis” (ISMARA) tool. ISMARA predicts regulatory interactions between the TFs and the target genes based on TF binding motifs [11]. For the SKCM data from TCGA only preprocessed data was available. Hence, ISMARA could not be used via the web portal. Therefore, the ISMARA analysis was performed based on the FPKM values (downloaded from the GDC portal [59], June 2018) by the developers using default settings.

## List of abbreviations

MIPRIP: Mixed Integer linear Programming based Regulatory Interaction Predictor
TMM: telomere maintenance mechanism
TF: transcription factor
GRN: gene regulatory network
ChIP: Chromatin immunoprecipitation
TCGA: The Cancer Genome Atlas
SKCM: skin cutaneous melanoma
OV: ovarian serous cystadenocarcinoma
CESC: cervical cancer
THYM: thymoma
TGCT: testicular germ cell cancer
PRAD: prostate adenocarcinoma
PAAD: pancreatic ductal adenocarcinoma
BRCA: breast cancer
ISMARA: Integrated Motif Activity Response Analysis
EMSA: electrophoretic shift assay
MILP: Mixed Integer Linear Programming
GDAC: Genome Data Analysis Center
RSEM: accurate transcript quantification from RNA-Seq data with or without a reference genome
ChEA: ChIP Enrichment Analysis
HTRIdb: Human Transcriptional Regulation Interactions database
TBA: total binding affinity
es: edge strength
FPKM: fragments per kilobase of exon model per million reads mapped
E-value: expected value
TERT: Telomerase reverse transcriptase
AR: androgen receptor
ETS: E-twenty six

## Declarations

### Acknowledgements

We thank the members of the CancerTelSys consortium for fruitful discussions on telomere maintenance mechanisms in cancer and Mikhail Pachkov for the ISMARA analysis. We further would like to thank Ashwini Kumar Sharma for his instructions how to download TCGA data and proofreading of the manuscript.

### Availability of data and material

The R-package ‘MIPRIP2’ and the generic human/mouse regulatory networks can be downloaded from https://www.leibniz-hki.de/en/miprip.html or https://github.com/network-modeling/MIPRIP. All datasets analyzed in this study are available at the GDAC (http://gdac.broadinstitute.org/) or the GDC (https://portal.gdc.cancer.gov/) portal.

### Authors’ contributions

PAM, OM, KT developed the Mixed Integer linear model, PAM implemented the MIPRIP 2.0 tool and performed the analysis. KT, AV and KA constructed the human and mouse generic gene regulatory network. PAM and KR wrote the manuscript with contributions from RK. All authors read and approved the final manuscript.

### Ethics approval and consent to participate

Not applicable.

### Consent for publication

Not applicable.

## References

1. Wright WE, Tesmer VM, Huffman KE, Levene SD, Shay JW: Normal human chromosomes have long G-rich telomeric overhangs at one end. Genes Dev 1997, 11(21):2801–2809.

2. Gaspar TB, Sa A, Lopes JM, Sobrinho-Simoes M, Soares P, Vinagre J: Telomere Maintenance Mechanisms in Cancer. Genes (Basel) 2018, 9(5).

3. Artandi SE, DePinho RA: Telomeres and telomerase in cancer. Carcinogenesis 2010, 31(1):9– 18.

4. Shay JW, Bacchetti S: A survey of telomerase activity in human cancer. Eur J Cancer 1997, 33(5):787–791.

5. Sandin S, Rhodes D: Telomerase structure. Curr Opin Struct Biol 2014, 25:104–110.

6. Feng J, Funk WD, Wang SS, Weinrich SL, Avilion AA, Chiu CP, Adams RR, Chang E, Allsopp RC, Yu J et al: The RNA component of human telomerase. Science 1995, 269(5228):1236–1241.

7. Kim NW, Piatyszek MA, Prowse KR, Harley CB, West MD, Ho PL, Coviello GM, Wright WE, Weinrich SL, Shay JW: Specific association of human telomerase activity with immortal cells and cancer. Science 1994, 266(5193):2011–2015.

8. Ramlee MK, Wang J, Toh WX, Li S: Transcription Regulation of the Human Telomerase Reverse Transcriptase (hTERT) Gene. Genes (Basel) 2016, 7(8):50.

9. Vinagre J, Almeida A, Populo H, Batista R, Lyra J, Pinto V, Coelho R, Celestino R, Prazeres H, Lima L et al: Frequency of TERT promoter mutations in human cancers. Nat Commun 2013, 4:2185.

10. Poos AM, Maicher A, Dieckmann AK, Oswald M, Eils R, Kupiec M, Luke B, Konig R: Mixed Integer Linear Programming based machine learning approach identifies regulators of telomerase in yeast. Nucleic Acids Res 2016, 44(10):e93.

11. Balwierz PJ, Pachkov M, Arnold P, Gruber AJ, Zavolan M, van Nimwegen E: ISMARA: automated modeling of genomic signals as a democracy of regulatory motifs. Genome Res 2014, 24(5):869–884.

12. Jiang P, Freedman ML, Liu JS, Liu XS: Inference of transcriptional regulation in cancers. Proc Natl Acad Sci U S A 2015, 112(25):7731–7736.

13. Li Y, Liang M, Zhang Z: Regression analysis of combined gene expression regulation in acute myeloid leukemia. PLoS Comput Biol 2014, 10(10):e1003908.

14. Frohlich H: biRte: Bayesian inference of context-specific regulator activities and transcriptional networks. Bioinformatics 2015, 31(20):3290–3298.

15. Lachmann A, Giorgi FM, Lopez G, Califano A: ARACNe-AP: gene network reverse engineering through adaptive partitioning inference of mutual information. Bioinformatics 2016, 32(14):2233–2235.

16. Alvarez MJ, Shen Y, Giorgi FM, Lachmann A, Ding BB, Ye BH, Califano A: Functional characterization of somatic mutations in cancer using network-based inference of protein activity. Nat Genet 2016, 48(8):838–847.

17. Schacht T, Oswald M, Eils R, Eichmuller SB, Konig R: Estimating the activity of transcription factors by the effect on their target genes. Bioinformatics 2014, 30(17):i401–407.

18. Horn S, Figl A, Rachakonda PS, Fischer C, Sucker A, Gast A, Kadel S, Moll I, Nagore E, Hemminki K et al: TERT promoter mutations in familial and sporadic melanoma. Science 2013, 339(6122):959–961.

19. Huang FW, Hodis E, Xu MJ, Kryukov GV, Chin L, Garraway LA: Highly recurrent TERT promoter mutations in human melanoma. Science 2013, 339(6122):957–959.

20. Wang L, Hurley DG, Watkins W, Araki H, Tamada Y, Muthukaruppan A, Ranjard L, Derkac E, Imoto S, Miyano S et al: Cell cycle gene networks are associated with melanoma prognosis. PLoS One 2012, 7(4):e34247.

21. R: A Language and Environment for Statistical Computing [https://www.R-project.org/]

22. MIPRIP website [https://www.leibniz-hki.de/en/miprip.html]

23. MIPRIP github repository [https://github.com/network-modeling/MIPRIP]

24. Babu MM, Luscombe NM, Aravind L, Gerstein M, Teichmann SA: Structure and evolution of transcriptional regulatory networks. Curr Opin Struct Biol 2004, 14(3):283–291.

25. Poole CJ, van Riggelen J: MYC-Master Regulator of the Cancer Epigenome and Transcriptome. Genes (Basel) 2017, 8(5).

26. Weintraub AS, Li CH, Zamudio AV, Sigova AA, Hannett NM, Day DS, Abraham BJ, Cohen MA, Nabet B, Buckley DL et al: YY1 Is a Structural Regulator of Enhancer-Promoter Loops. Cell 2017, 171(7):1573–1588 e1528.

27. Bougel S, Renaud S, Braunschweig R, Loukinov D, Morse HC, 3rd, Bosman FT, Lobanenkov V, Benhattar J: PAX5 activates the transcription of the human telomerase reverse transcriptase gene in B cells. J Pathol 2010, 220(1):87–96.

28. Chen YJ, Campbell HG, Wiles AK, Eccles MR, Reddel RR, Braithwaite AW, Royds JA: PAX8 regulates telomerase reverse transcriptase and telomerase RNA component in glioma. Cancer Res 2008, 68(14):5724–5732.

29. Moehren U, Papaioannou M, Reeb CA, Grasselli A, Nanni S, Asim M, Roell D, Prade I, Farsetti A, Baniahmad A: Wild-type but not mutant androgen receptor inhibits expression of the hTERT telomerase subunit: a novel role of AR mutation for prostate cancer development. FASEB J 2008, 22(4):1258–1267.

30. Crowe DL, Nguyen DC, Tsang KJ, Kyo S: E2F-1 represses transcription of the human telomerase reverse transcriptase gene. Nucleic Acids Res 2001, 29(13):2789–2794.

31. Chebel A, Ffrench M: Transcriptional regulation of the human telomerase reverse transcriptase: new insights. Transcription 2010, 1(1):27–31.

32. Mani KM, Lefebvre C, Wang K, Lim WK, Basso K, Dalla-Favera R, Califano A: A systems biology approach to prediction of oncogenes and molecular perturbation targets in B-cell lymphomas. Mol Syst Biol 2008, 4:169.

33. Li Y, Zhou QL, Sun W, Chandrasekharan P, Cheng HS, Ying Z, Lakshmanan M, Raju A, Tenen DG, Cheng SY et al: Non-canonical NF-kappaB signalling and ETS1/2 cooperatively drive C250T mutant TERT promoter activation. Nat Cell Biol 2015, 17(10):1327–1338.

34. Rouaud F, Hamouda-Tekaya N, Cerezo M, Abbe P, Zangari J, Hofman V, Ohanna M, Mograbi B, El-Hachem N, Benfodda Z et al: E2F1 inhibition mediates cell death of metastatic melanoma. Cell Death Dis 2018, 9(5):527.

35. Chiappetta G, Avantaggiato V, Visconti R, Fedele M, Battista S, Trapasso F, Merciai BM, Fidanza V, Giancotti V, Santoro M et al: High level expression of the HMGI (Y) gene during embryonic development. Oncogene 1996, 13(11):2439–2446.

36. Agostini A, Panagopoulos I, Andersen HK, Johannesen LE, Davidson B, Trope CG, Heim S, Micci F: HMGA2 expression pattern and TERT mutations in tumors of the vulva. Oncology reports 2015, 33(6):2675–2680.

37. Griewank KG, Murali R, Puig-Butille JA, Schilling B, Livingstone E, Potrony M, Carrera C, Schimming T, Moller I, Schwamborn M et al: TERT promoter mutation status as an independent prognostic factor in cutaneous melanoma. J Natl Cancer Inst 2014, 106(9).

38. Firehose stddata 2016_01_28 run [http://gdac.broadinstitute.org/]

39. Publication Guidlines [http://cancergenome.nih.gov/publications/publicationguidelines]

40. Li B, Dewey CN: RSEM: accurate transcript quantification from RNA-Seq data with or without a reference genome. BMC Bioinformatics 2011, 12:323.

41. MetaCore [https://portal.genego.com/]

42. Lachmann A, Xu H, Krishnan J, Berger SI, Mazloom AR, Ma’ayan A: ChEA: transcription factor regulation inferred from integrating genome-wide ChIP-X experiments. Bioinformatics 2010, 26(19):2438–2444.

43. Chen L, Wu G, Ji H: hmChIP: a database and web server for exploring publicly available human and mouse ChIP-seq and ChIP-chip data. Bioinformatics 2011, 27(10):1447–1448.

44. Bovolenta LA, Acencio ML, Lemke N: HTRIdb: an open-access database for experimentally verified human transcriptional regulation interactions. BMC Genomics 2012, 13:405.

45. Yang JH, Li JH, Jiang S, Zhou H, Qu LH: ChIPBase: a database for decoding the transcriptional regulation of long non-coding RNA and microRNA genes from ChIP-Seq data. Nucleic Acids Res 2013, 41(Database issue):D177-187.

46. Grassi E, Zapparoli E, Molineris I, Provero P: Total Binding Affinity Profiles of Regulatory Regions Predict Transcription Factor Binding and Gene Expression in Human Cells. PLoS One 2015, 10(11):e0143627.

47. Loots G, Ovcharenko I: ECRbase: database of evolutionary conserved regions, promoters, and transcription factor binding sites in vertebrate genomes. Bioinformatics 2007, 23(1):122–124.

48. Essaghir A, Toffalini F, Knoops L, Kallin A, van Helden J, Demoulin JB: Transcription factor regulation can be accurately predicted from the presence of target gene signatures in microarray gene expression data. Nucleic Acids Res 2010, 38(11):e120.

49. Zhao F, Xuan Z, Liu L, Zhang MQ: TRED: a Transcriptional Regulatory Element Database and a platform for in silico gene regulation studies. Nucleic Acids Res 2005, 33(Database issue):D103–107.

50. Kel AE, Kolchanov NA, Kel OV, Romashchenko AG, Anan’ko EA, Ignat’eva EV, Merkulova TI, Podkolodnaia OA, Stepanenko IL, Kochetov AV et al: [TRRD: a database of transcription regulatory regions in eukaryotic genes]. Mol Biol (Mosk) 1997, 31(4):626–636.

51. Portales-Casamar E, Kirov S, Lim J, Lithwick S, Swanson MI, Ticoll A, Snoddy J, Wasserman WW: PAZAR: a framework for collection and dissemination of cis-regulatory sequence annotation. Genome Biol 2007, 8(10):R207.

52. Gronostajski RM, Guaneri J, Lee DH, Gallo SM: The NFI-Regulome Database: A tool for annotation and analysis of control regions of genes regulated by Nuclear Factor I transcription factors. J Clin Bioinforma 2011, 1(1):4.

53. Gurobi Optimizer Reference Manual [http://www.gurobi.com]

54. Cancer Genome Atlas N: Genomic Classification of Cutaneous Melanoma. Cell 2015, 161(7):1681–1696.

55. Benjamini Y, Hochberg Y: Controlling the false discovery rate: a practical and powerful approach to multiple testing. J Roy Statist Soc Ser B 1995, 57:289–300.

56. Breitling R, Armengaud P, Amtmann A, Herzyk P: Rank products: a simple, yet powerful, new method to detect differentially regulated genes in replicated microarray experiments. FEBS Lett 2004, 573(1-3):83–92.

57. PubMed: 2018.

58. Smedley D, Haider S, Durinck S, Pandini L, Provero P, Allen J, Arnaiz O, Awedh MH, Baldock R, Barbiera G et al: The BioMart community portal: an innovative alternative to large, centralized data repositories. Nucleic Acids Res 2015, 43(W1):W589-598.

59. GDC Data Portal [https://portal.gdc.cancer.gov/]

