## Supplementary_material for "Modelling *TERT* regulation across 19 different cancer types based on the MIPRIP 2.0 gene regulatory network approach"

**Table S1: Selected cancers from The Cancer Genome Atlas**

| Cancer type | Number of tumor samples |
| --- | --- |
| Breast cancer (BRCA) | 983 |
| Cervical cancer (CESC) | 300 |
| Colorectal adenocarcinoma (COADREAD) | 619 |
| Cutaneous melanoma (SKCM) | 103 |
| Glioblastoma multiforme (GBM) | 145 |
| Head and neck squamous cell carcinoma (HNSC) | 501 |
| Liver hepatocellular carcinoma (LIHC) | 342 |
| Lung adenocarcinoma (LUAD) | 491 |
| Lung squamous cell carcinoma (LUSC) | 489 |
| Ovarian serous cystadenocarcinoma (OV) | 294 |
| Prostate adenocarcinoma (PRAD) | 445 |
| Stomach adenocarcinoma (STAD) | 405 |
| Urothelial bladder cancer (BLCA) | 399 |
| Uterine corpus endometrial carcinoma (UCEC) | 532 |
| Acute Myeloid Leukemia (LAML) | 168 |
| Testicular germ cell cancer (TGCT) | 148 |
| Esophageal cancer (ESCA) | 178 |
| Pancreatic ductal adenocarcinoma (PAAD) | 145 |
| Thymoma (THYM) | 120 |

**Table S2: List of transcription factors putatively regulating *TERT*, from the generic human gene regulatory network**

| TF | Edge strength score | TF | Edge strength score | TF | Edge strength score |
|---|---|---|---|---|---|
| **AP-2** | 2.00 | **HMGA2** | 2.00 | **PITX1** | 2.00 |
| **AR** | 2.00 | **HNRNPK** | 2.00 | **POLR2A** | 0.50 |
| **BATF** | 0.25 | **IKZF1** | 2.00 | **POU2F2** | 0.25 |
| **BCL11A** | 0.25 | **IRF1** | 2.00 | **RAD21** | 0.50 |
| **BHLHE40** | 2.00 | **JUND** | 2.00 | **RELA** | 2.00 |
| **CEBPA** | 2.00 | **KLF2** | 2.00 | **REST** | 0.25 |
| **CTCF** | 0.50 | **MAX** | 2.50 | **RUNX2** | 2.00 |
| **CTCFL** | 2.00 | **MAZ** | 2.00 | **SIN3A** | 0.50 |
| **E2F1** | 2.00 | **MEN1** | 2.00 | **SIN3AK20** | 0.50 |
| **E2F2** | 2.00 | **MITF** | 2.00 | **SMAD3** | 2.00 |
| **E2F4** | 2.25 | **MXD1** | 2.00 | **SMARCB1** | 0.25 |
| **E2F5** | 2.00 | **MXI1** | 0.50 | **SP3** | 2.00 |
| **E2F6** | 2.25 | **MYB** | 2.00 | **TAF1** | 0.25 |
| **EGR1** | 2.50 | **MYC** | 3.75 | **TAF9** | 2.00 |
| **EPAS1** | 2.00 | **MYCN** | 2.00 | **TAL1** | 2.00 |
| **ESR1** | 2.00 | **MZF1** | 2.00 | **TCF12** | 0.25 |
| **ESR2** | 2.00 | **NFAT5** | 2.00 | **TCF7** | 2.00 |
| **ETS1** | 2.00 | **NFATC2** | 2.00 | **TFAP2A** | 2.00 |
| **ETS2** | 2.00 | **NF.KB** | 2.00 | **TFAP2B** | 2.00 |
| **GLI1** | 2.00 | **NFKB1** | 1.00 | **TFAP2C** | 2.00 |
| **GLI2** | 2.00 | **NFKB.P50.P65** | 2.00 | **TFAP2D** | 2.00 |
| **GRHL2** | 2.00 | **NFX1** | 2.00 | **TP53** | 2.00 |
| **HEY1** | 0.25 | **NR2F2** | 2.00 | **TP73** | 2.00 |
| **HIF.1** | 2.00 | **PAX5** | 2.00 | **WT1** | 2.00 |
| **HIF1A** | 2.00 | **PAX8** | 2.00 | **ZBTB48** | 2.00 |

**Table S3: Specific *TERT* regulators of each cancer type (see Excel-file)**

### Table S4: Number of Pubmed hits for the predicted common *TERT* regulators

| Regulator | Pubmed hits with *TERT* (AND “telomerase” AND “human” AND “regulation”) | Pubmed hits without *TERT* (AND ”telomerase” AND “human” AND “regulation”) |
|---|---|---|
| **E2F4** | 0 | 3 |
| **AR** | 15 | 35 |
| **PAX5** | 2 | 2 |
| **E2F2** | 1 | 2 |
| **BATF** | 2 | 2 |
| **PAX8** | 1 | 3 |
| **SMARCB1** | 0 | 0 |
| **MXI1** | 0 | 0 |
| **TAF1** | 0 | 0 |

### Table S5: Confusion matrix for the Pubmed query)

| | Found with the query containing the nine predicted regulators | Found only with the query which did not contain the nine predicted regulators |
|---|---|---|
| **Query with TERT** | 21 | 981 |
| **Query w/o TERT** | 25 | 2,483 |

**Table S6: *TERT* regulators predicted with ISMARA**

| Regulator | Score |
|---|---|
| AHR_ARNT2 | 0.38 |
| ARNT | 0.73 |
| CTCF_CTCFL | 0.85 |
| ELF2_GABPA_ELF5 | 0.01 |
| ELK4_ETV5_ELK1_ELK3_ELF4 | 1.41 |
| GMEB2 | 1.11 |
| HES1 | 0.33 |
| IKZF1 | 0.23 |
| KLF16_SP2 | 1.48 |
| MAZ_ZNF281_GTF2F1 | 0.41 |
| MNT_HEY1_HEY2 | 1.05 |
| MXI1_MYC_MYCN | 3.40 |
| MYF6 | 0.10 |
| PLAGL1 | 1.37 |
| RCOR1_MTA3 | 0.85 |
| SIN3A_CHD1 | 0.84 |
| SIX4 | 1.31 |
| TCF3_MYOG | 0.32 |
| TCF12_ASCL2 | 1.05 |
| WT1_MTF1_ZBTB7B | 0.04 |
